# Comet fragment-ion indexing for enhanced peptide sequencing

**DOI:** 10.1101/2024.10.11.617953

**Authors:** Christopher D. McGann, Erik J. Bergstrom, Vagisha Sharma, Lilian R. Heil, Qing Yu, Jimmy K. Eng, Devin K. Schweppe

## Abstract

Fragment ion indexing has significantly improved the efficiency of proteomics database search tools. This work implements fragment ion indexing in Comet, a widely-used, open-source search engine. We demonstrate that this enhancement maintains scoring and identification accuracy while substantially increasing peptide spectral matching speeds across multiple applications, including open modification searches, immunopeptidomics, and real-time searches. Comet-FI reduced search speeds by up to 94%, enabling the rapid analysis of complex data types like immunopeptidomes. Fragment-ion indexing enables Comet to keep pace with modern instrumentation and expanding applications in proteomics, reinforcing its utility in diverse proteomics workflows and its integration with a wide range of proteomics tools and platforms.

## Main Text

Mass spectrometry is capable of producing thousands of tandem mass spectra per minute^1^. Database searching has remained the most common method of identifying peptides from these acquired spectra. Scoring each experimental spectrum against a database of in-silico generated theoretical peptides has proven to be a robust and sensitive approach that has improved over time^2^. Descending from the original database search algorithm, Comet, has remained one of the most popular search tools for over a decade^3–5^. Comet’s free and open-source nature has led to its inclusion in a myriad of proteomics workflows and projects^6–15^, establishing itself as a hallmark of the field. Comet has been both a critical tool for individual users and a key component of several widely-used proteomics platforms, such as quantms^16^, Galaxy^17^, and OpenMS^7^. As the field of mass spectrometry proteomics has expanded, so has the application of database searching. With this expansion comes the need for tools to handle the latest developments, driving ever increasing numbers of samples and spectra that must be analyzed. Techniques such as open-modification searching, non-specific immunopeptidomic searches, proteogenomic searching, and real-time searching are exciting but impose significant computational demands. Alongside the ongoing advancements in instrumentation, this creates a pertinent need to improve the efficiency of database peptide spectral matching.

Fragment ion indexing has been a major advancement in speeding up the search process^18^. Originally implemented by MSFragger, this inverted index style approach involves in-silico digestion of proteins into a database of theoretical fragment ions, allowing the filtering of candidate peptides based on the presence of these ions (in addition to filtering based on the precursor tolerance). Fragment ion indexing significantly reduces spectral scoring time and has been effectively applied to tasks such as open-modification searches^19^. This strategy has been adopted by other tools as well, achieving impressive search speeds^20^. In this study, we adapt a fragment ion indexed approach for the widely-used, open-source Comet search algorithm, significantly accelerating its search capabilities (Fig. 1A). This enhancement enables Comet to keep pace with new applications and modern instrumentation, maintaining its relevance as a key tool in proteomics research.

**Figure 1.**
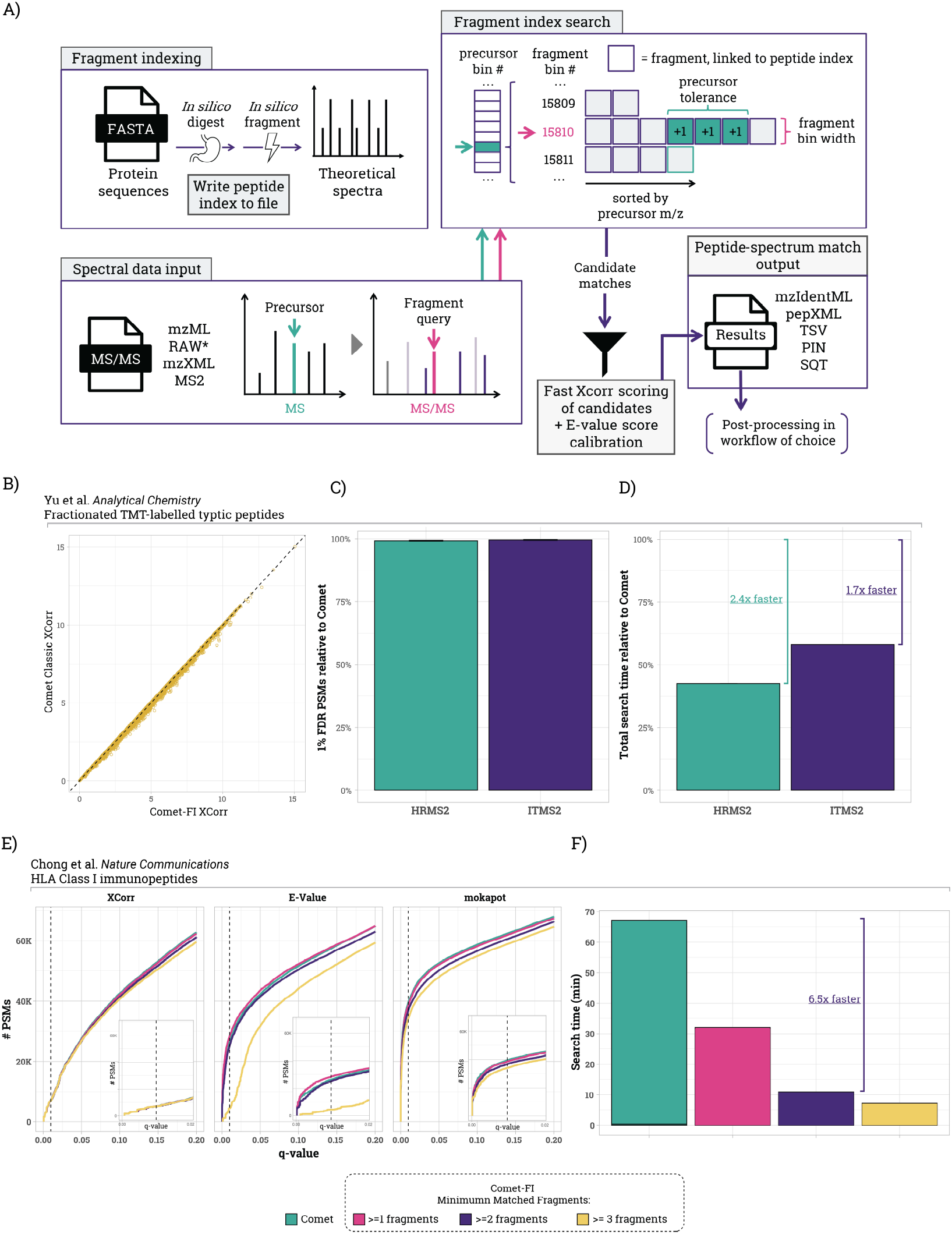
Overview of fragment indexing in Comet. A) Graphical workflow describing the new indexing strategy integrated into Comet. B) Comparison of XCorr values between Comet and Comet-FI for the same peptide spectral match. C) Number of FDR filtered peptide spectral matches for TMT-labeled fractionated data using Comet-FI as compared to baseline with Comet. D)Total search time relative to Comet baseline for same dataset E) PSMs at different q-values as determined by XCorr, E-value, or mokapot scores for HLA Class I data. The dotted line indicates a q-value of 0.01 (or 1%). Inset plots zoom in on the q-value range from 0 to 0.02. Classic search is precursor indexed and has no minimum matched fragments. F) Search times for the raw files shown in E.

The main cross correlation (XCorr) scoring function, which is well-established in the field^2,3,21^, was maintained moving from Comet to Comet-FI. As such, comparison of XCorr values for peptide-spectral matches (PSMs) from the same analytical run for Comet and Comet-FI resulted in near-identical values (Fig.1B). Extending this to a twelve fraction TMT-labeled dataset, we detect a very similar number of filtered PSMs for both linear ion trap and Orbitrap spectra at a 1% FDR as calculated by mokapot. Notably, using Comet-FI, these searches finished 1.7-fold faster for the ITMS2 data and 2.3-fold faster for the HRMS2 data with no significant change in total filtered PSMs (Fig. 1C, 1D).

Comet calculates a calibrated score called the expectation value (E-value) which is better for target-decoy discrimination and FDR estimation^4^. The E-value is calculated based on fitting a linear regression to the survival function of the XCorr score distribution for a given spectrum^22^. Fragment ion indexing allows for candidate peptides with matched fragment counts below a threshold to not be scored. We’ve observed that despite the low chance that a peptide with 1 to 2 matched fragments is a correct identification, skipping these peptides alters the XCorr score distributions and can affect the discriminating power of the E-value (Fig. 1E). Peptidomic studies often require searches unconstrained by enzyme specificity, where all possible unique peptides within a given length range (e.g., 8-15 amino acids) are considered. The large pool of candidate peptides provides a good dataset for evaluating the relationship between the minimum number of fragment ions considered, the E-values reported, and the total sensitivity for PSM detection. Benchmarking on HLA-I data with challengingly large search spaces due to non-enzymatic digestions, we evaluated how changing the minimum number of fragment ions required for scoring affects the number of peptide spectral matches across the range of q-values using XCorr, E-value, or mokapot’s discriminant score to rank PSMs. Using E-value alone with two or more fragment ions severely reduced the number of PSMs passing FDR filters. (Fig. 1E). As such, it is inadvisable to use an E-value in isolation for FDR determination. However, when used with programs like mokapot or Percolator, there is very little difference between 0, 1, and 2 matched ions. Using the default parameters with a minimum of two matched fragments, Comet-FI reduced the runtime of HLA peptide searches by 6.5-fold (Fig. 1F).

The latest generation of mass spectrometers can acquire spectra at 300 Hz, producing data at a rate that makes post-hoc peptide identification one of the rate limiting steps in the complete analysis workflow^23^. To test Comet-FI on these modern instruments with a large and challenging search space, we analyzed data from Guzman et al. 2024 (PXD046453)^24,25^ collected with a wide 2 Th precursor isolation window DDA method on the Orbitrap Astral. Based on the isolation widths, we used Comet-FI with a precursor tolerance of ±1 Th (vs the default Comet tolerance of ± 20 ppm). At 1% FDR, we observed a 3.6% boost in identified PSMs for Comet-FI compared to Comet (Fig. 2A). More importantly, Comet-FI was 3.43-fold faster than Comet when searching these Astral data (Comet 155.6 min; Comet-FI 45.3 min) (Fig. 2B). Comet-FI’s enhanced search speed means that post hoc searching can now keep up with the increasing speed of modern, high-speed instruments.

**Figure 2.**
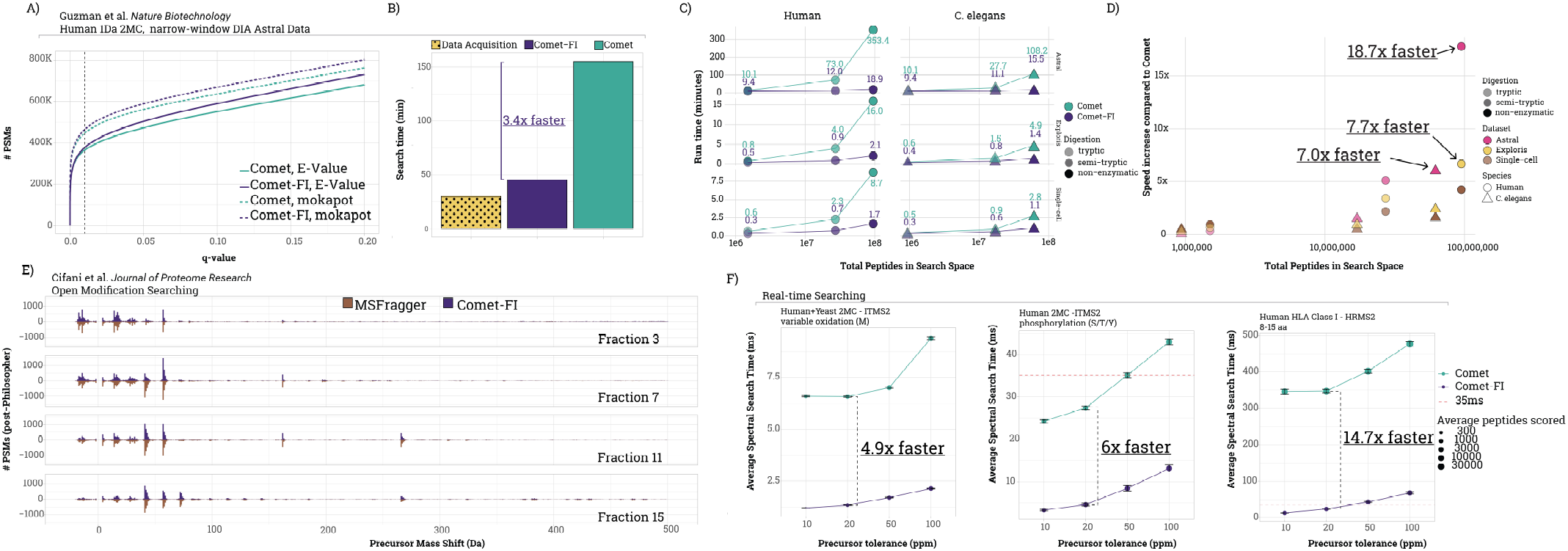
Evaluating Comet-FI in various workflows. A) Peptide spectral matches at different q-values using either E-value or mokapot score for Comet and Comet-FI using narrow window DIA data collected on an Astral. B) Total search times for the same data using Comet and Comet-FI with the instrument acquisition time also plotted. C) Relationship between run time and total peptides in search space of different datasets using tryptic, semi-tryptic, and non-enzymatic digestion settings (see Table S1 for file names). D) The relative increase of single file search speed increases compared to classic Comet from the data portrayed in C. E) Precursor mass shift on peptide spectral matches from running Philosopher with Comet-FI or MSFragger. -1.5 to 3.5 Da are excluded from the plot for readability. Fractions 3, 7, 11, and 15 are plotted. F) Real-time search time comparisons for various data types and search parameters as described above each plot. Red dotted line at 35 ms is the standard MS2 fill time as part of an RTS-MS3 based workflow^34^. Data in F were collected in this study (left and middle panels) or Stopfer et al. (PXD029860, right panel)^35^.

To explore the relationship between search speed and large search spaces due to database size and enzymatic constraints, we queried one file of the TMT-labeled dataset against yeast and human databases using three different enzyme constraints (fully tryptic, semi-tryptic, and non-enzymatic). For this analysis, we compared both the total number of peptides as the total search space and the number of PSMs that were eventually scored by the XCorr calculation. As shown in Figure 2C, the search space grows significantly as the enzyme constraint is relaxed. Varying the *in silico* digestion shows that constraints applied during fragment-ion indexing (e.g., number of required fragments) greatly reduces the number of PSMs that are eventually scored compared to classic Comet. In the case of human database searches, there is a ∼400 to ∼800-fold reduction in PSMs that are scored by XCorr when fragment-ion indexing is applied. The result of the reduced scoring burden was an 18-fold increase in search speeds for non-enzymatic searches with Comet-FI compared to classic Comet (Table S1, Fig. 2C, 2D).

Searches with a very wide precursor tolerance (hundreds of daltons) have been used to identify genetic and chemical variants of canonical peptide sequences^19,26,27^. Open searching dramatically increases the candidate peptide sequence space, necessitating improved search speeds ^18,20^. Thus, while Comet has been used for open search pipelines^28^, the compute time was not practical on common laboratory workstations. To evaluate the utility of Comet-FI for open searching, we used the Philosopher software pipeline which includes both Comet and the closed-source fragment indexing search engine, MSFragger^18^. Comet-FI for open searching was able to generate data consistent with MSFragger’s established workflow (Fig. 2E). Importantly, Comet-FI performed open-searching ∼50-times faster than Comet. Notably, Comet’s search times were previously reported to be over 151-times slower than MSFragger. While MSFragger’s closed-source search algorithm was still faster than Comet-FI, the difference in speeds for open-searching was reduced to 3.8-fold (Fig. S1, average of 21 min for MSFragger and 80 min per file for Comet-FI). Thus, Comet-FI’s open-source fragment ion indexing implementation can readily be used for open searching and easily integrated anywhere that Comet has classically been used before. We note that MSFragger has been actively developed since 2017. Thus, ongoing Comet-FI code development, optimizations, and improvements are on track to offer a premier, open-source alternative to other fragment-ion indexing approaches.

Real-time searching (RTS), i.e., simultaneous identification of peptides with instrument acquisition, allows mass spectrometric methods to use peptide sequence information to inform spectral acquisition^15,29–32^ with applications to isobarically multiplexed samples (TMT-labeled) and single-cell proteomics^33^. Owing to the open-source codebase, Comet was the first full-featured algorithm used for TMT-based RTS methods and the first RTS integrated into vendor instrument control software^34^. While Comet’s classic implementation worked well for canonical proteomics applications (single species database, unmodified peptide sequences, minimal missed cleavages, sub-50 ppm precursor tolerances, etc.), for applications requiring larger sequence search spaces, the spectral search time often exceeded the MS2 acquisition time (35 ms). Therefore, it could not be perfectly parallelized with instrument acquisition. This was particularly true for searches with multiple missed cleavages, wide precursor windows, variable modifications (e.g. phosphoproteomics), and immunopeptidomics searches for HLA-bound peptides. Comet-FI integrated in the Orbiter real-time search platform^15^ reduced the median single-spectrum search times for RTS by 4.9-fold for tryptic peptide searches, 6-fold for phosphopeptide searches, and 14.7-fold for HLA Class I peptide searches at 20 ppm precursor tolerance (Fig. 2G, S2). Single spectrum search times that are commensurate with MS2 acquisition rates for both unmodified and modified searches, as well as HLA-I derived peptides, are now possible with Comet-FI for RTS, opening the door for biologically-aware instrument control.

The implementation of fragment ion indexing in the open-source Comet search algorithm significantly enhances post hoc and real-time spectral searching for proteomics analyses across data and instrument types. Comet-FI accelerates search times, enables practical open searching, and increases the real-time analysis capabilities with adaptable and extensible code based on well-established scoring and spectral matching. Comet’s open-source code, ecosystem of developers, and track record of diverse implementations will ensure that the enhanced performance and versatility of Comet-FI will provide ongoing benefit for the proteomics research community. These advancements position the Comet search engine to keep pace with the ongoing evolution of the proteomics field.

## Data Availability

All new data has been deposited at: MSV000096008.

## Acknowledgements

The authors would like to thank all Comet contributors and users, especially Drs. WIll Fondrie and Michael Hoopmann. We thank Drs. Jesse Canterbury, Graeme McAlister, Mike Senko, and Will Barshop for technical discussions. We thank Ethan Crawford for technical assistance in code development. We would also like to acknowledge our funding sources: R35GM150919-01 (DKS), Andy Hill CARE Distinguished Researcher Award (DKS), Cancer Consortium New Investigator Award (P30 CA015704, DKS), the Pew Charitable Trusts (DKS).

## Online Methods

### Fragment ion indexing in Comet

MS/MS spectra are pre-processed by representing peaks in an 1D array, binning masses based on the fragment bin tolerance/offset parameters. To generate the fragment ion index, Comet first generates a peptide index file, denoted with an “.idx” file name extension, based on search/digest parameters (Figure 1A). To perform a fragment ion index search, the user specifies the peptide index file instead of a standard FASTA as the query database. The peptide index contains peptide sequences, their unmodified masses, file positions of each parent protein within the original protein sequence FASTA database, and combinatorial bitmasks representing potential variable modification positions. At search time, a user controlled setting can direct Comet to calculate fragment ion m/z (y3-yn, b3-bn) only for theoretical peptides with precursor m/z’s detected in the input spectrum and bins fragments by the same tolerance used to pre-process spectra. The fragment ion index is segmented per search thread. And to mitigate over-populating any specific index entry, the index is further divided into a large number of precursor m/z bins. Binned fragments are then queried against the fragment ion index and the number of matched ions for any potential peptide hits are incremented. Candidate peptides are filtered based on the number of matched ions before undergoing full XCorr^21^ scoring and E-value^22^ calculation. Output files contain the same content as a non-indexed search and are compatible with the same downstream workflows.

### Analysis of publicly available data

To evaluate the performance of Comet’s newly implemented indexing strategy, publicly available datasets (PXD016766^34^, PXD046453^25^, PXD013649^36^, PXD029860^35^, and PXD019853^37^) retrieved from

ProteomeXchange^38^ were searched with and without a fragment ion index. Searches were run on a desktop computer (Intel i9-11900) using 12 threads and 64 GB memory on Windows Subsystem for Linux. Peptide-spectrum matches (PSMs) were exported in the Percolator input file format and re-scored using mokapot^39^, a Python implementation of the widely used Percolator approach for target-decoy discrimination^40^. For false discovery rate estimation, mokapot ^39^ was used to aggregate multiple features into a single score, using a Support Vector Machine classifier, to increase the sensitivity of peptide detections. If applicable, protein inference was also performed using mokapot. PSM, peptide, and protein identifications using each search mode were evaluated using Comet’s XCorr score, E-value score, and the mokapot post-processing score^39^. In the Philosopher^41^ open search, the recommended workflow was used.

### Data collection for real-time search analyses

Data for some real-time search benchmarks was collected in house. Human+yeast and human phosphoproteomics samples were prepared as previously described^42–44^. For the human+yeast sample, peptides were eluted over 60 min gradients running from 96% Buffer A (5% acetonitrile, 0.125% formic acid) and 4% buffer B (95% acetonitrile, 0.125% formic acid) to 30% buffer B. Sample eluate was electrosprayed (2700 V) into a Thermo Scientific Orbitrap Eclipse mass spectrometer for analysis. High field asymmetric waveform ion mobility spectrometry (FAIMS) was set at “standard” resolution, 4.6 L/min gas flow, and 3 CVs: −40/–60/–80 were used. MS1 scans were conducted at 120,000 resolving power with a 50 ms max injection time, and the AGC target set to 100%. Peaks from the MS1 scans were filtered by intensity (minimum intensity >5 × 10^3^), charge state (2 ≤ z ≤ 6), and detection of a monoisotopic mass (monoisotopic precursor selection, MIPS). Dynamic exclusion was used with a duration of 60 s, repeat count of 1, mass tolerance of 10 ppm, and the “exclude isotopes” option checked. For each MS1, 8 data-dependent MS/MS scans were collected. MS/MS scans were conducted in the linear ion trap with the “rapid” scan rate, 50 ms max injection time, AGC target set to 200%, CID collision energy of 35% with 10 ms activation time, and 0.5 m/z isolation window. SPS ions were set to 10 and MS3 scans were performed at a resolving power of 50,000, with an HCD collision energy of 45%, AGC of 200%, with a maximum injection time of 200 ms.

Phosphopeptides were eluted over a 120 min gradient running from 94% Buffer A (5% acetonitrile, 0.125% formic acid) and 4% buffer B (80% acetonitrile, 0.125% formic acid) to 28% buffer B. Sample eluate was electrosprayed (2700 V) into a Thermo Scientific Orbitrap Ascend mass spectrometer for analysis. High field asymmetric waveform ion mobility spectrometry (FAIMS) was set at “standard” resolution, 4.6 L/min gas flow, and 3 CVs: −40/–60/–80 were used. MS1 scans were conducted at 120,000 resolving power with a 251 ms max injection time, and the AGC target set to 100%. Peaks from the MS1 scans were filtered by intensity (minimum intensity >5 × 10^3^), charge state (2 ≤ z ≤ 6), and detection of a monoisotopic mass (monoisotopic precursor selection, MIPS). Dynamic exclusion was used, with a duration of 90 s, repeat count of 1, mass tolerance of 10 ppm, and the “exclude isotopes” option checked. For each MS1, 8 data-dependent MS/MS scans were collected. MS/MS scans were conducted in the linear ion trap with the “rapid” scan rate, 75 ms max injection time, AGC target set to 250%, HCD normalized collision energy of 32% and 0.5 m/z isolation window. Phosphoproteomic real-time search database settings included a precursor tolerance of 20 ppm, max 2 missed cleavages, 7-50 amino acids in length, oxidation on M and phosphorylation on S/T/Y as variable modifications, maximum 3 variable mods per peptide. Immunopeptidomic real-time search database settings included a precursor tolerance of 20 ppm and a non-enzymatic digest of 8 to 15 amino acids long. Real-time search results were simulated using the publicly available real-time search option for Comet through in-house platform, Orbiter^15^.

All searches were performed against a target-decoy database where the decoy sequences were composed of reversing each protein sequence and appending those reverse sequences to the target sequence database. After ranking the Comet search results by either the XCorr, E-value, or mokapot^39^ score FDR was calculated using target and decoy peptides as 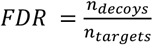 when using XCorr and E-value or 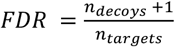 when using mokapot.

## Code Availability

Fragment ion indexed Comet is freely available under an Apache 2.0 license with version v2024.02.0 and newer on Github, https://github.com/UWPR/Comet. Classic search can be run the same as previous versions. A fragment ion index search is invoked by specifying a peptide index file instead of a FASTA file for the search database. Precursor search tolerance parameters were updated to allow asymmetric windows. Minimum number of matched fragment ions for reporting and scoring can be set with the “fragindex_min_ions_report” and “fragindex_min_ions_score” parameters, respectively.

