## Supplemental Figures for "Comet fragment-ion indexing for enhanced peptide sequencing"

### Supplemental Information

**Figure S1**. Open modification search timing.

**Figure S2**. Real time search timing boxplots.

**Figure S3.** Open modification real time search histogram.

A)

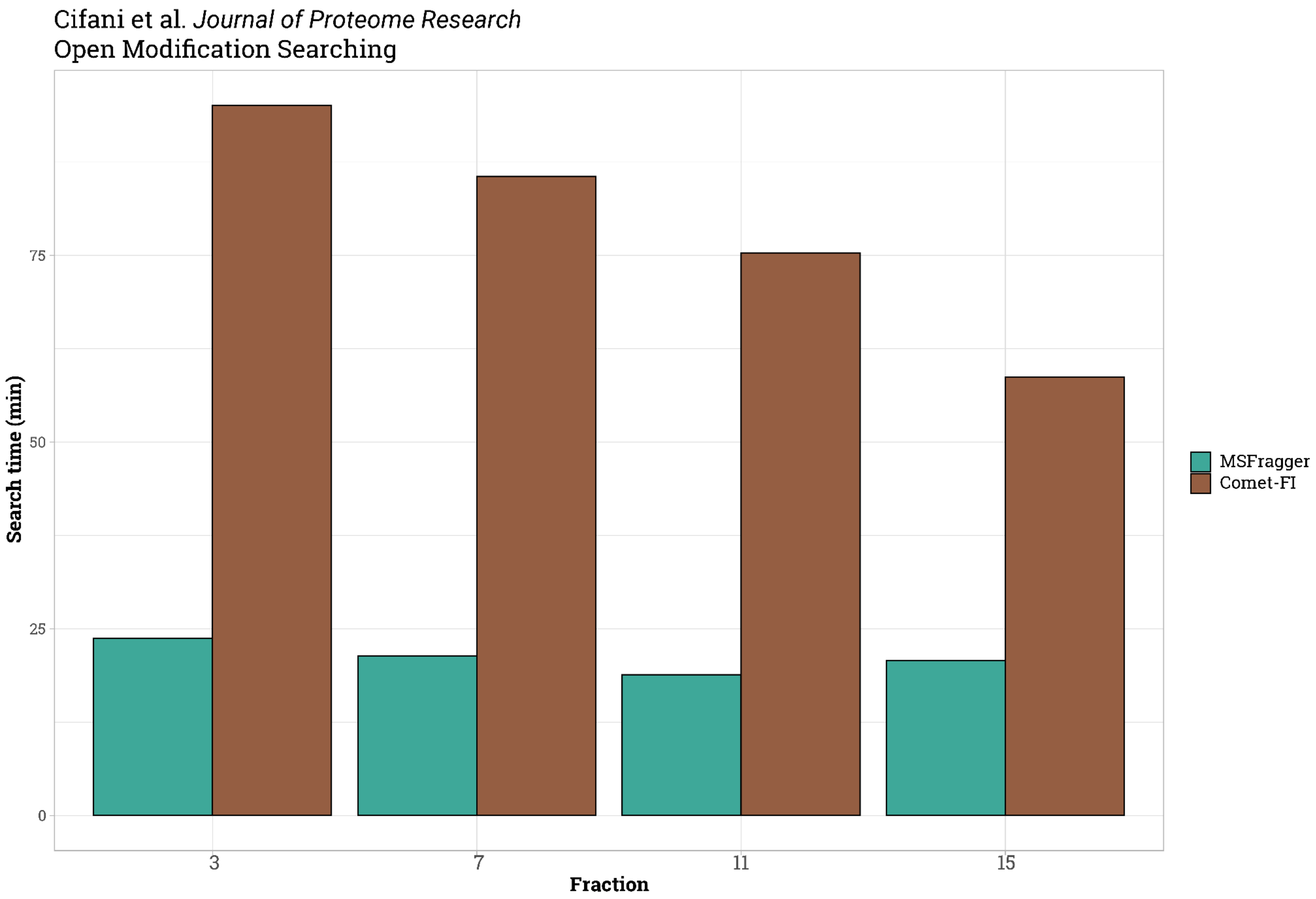


**Figure S1. Open Modification search timing.** A) Search time in minutes from running open search on fractions from Cifani et al. with MSFragger and Comet-FI.

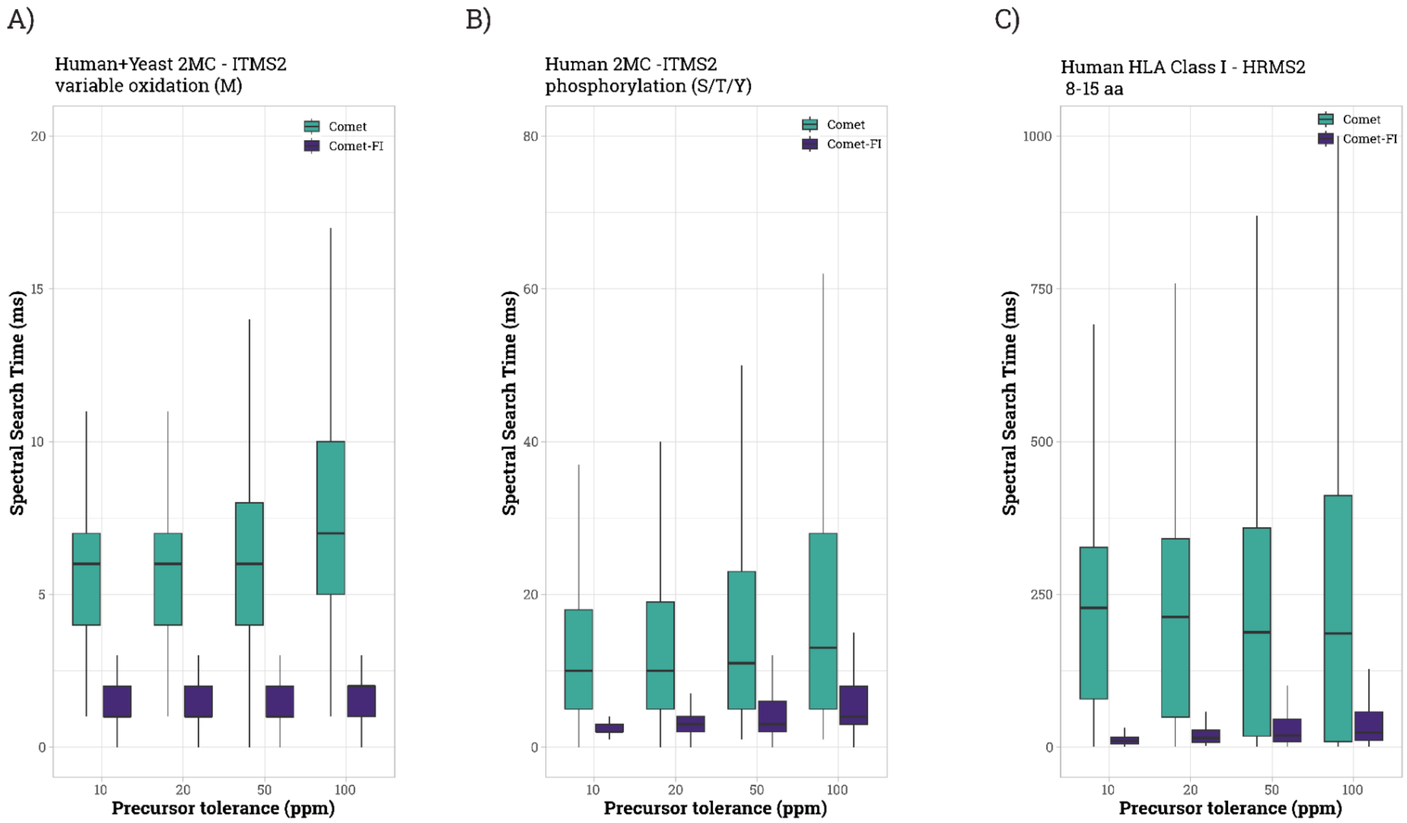


**Figure S2. Real time search.** A-C) Boxplots of RTS spectral search time for different search parameters.

A)

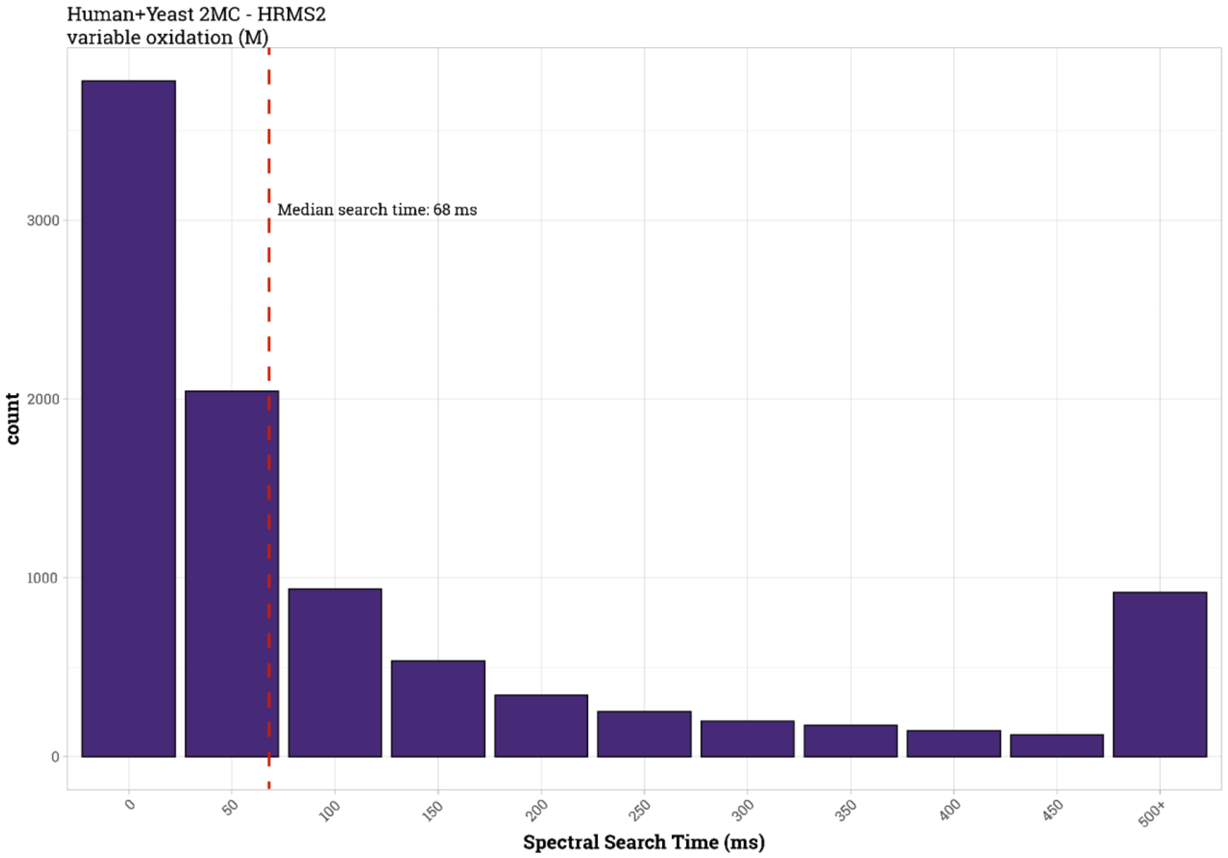


**Figure S3. Open Modification RTS.** Histogram of search times for open modification (-150 to +500 Da) real-time searching. Red dotted line is at the median search time.
